# First genome assemblies of Neotropical *Thoracobombus* Bumblebees *Bombus pauloensis* and *Bombus pullatus*

**DOI:** 10.1101/2025.10.13.682240

**Authors:** Andres Felipe Lizcano-Salas, Jesús Camilo Jacome-García, Diego Riaño-Jiménez, Marcela Guevara-Suarez

## Abstract

Bumble bees (*Bombus*) are considered to be essential pollinators of a wide range of flowering plants, within both agricultural and natural ecosystems. *Bombus pauloensis* and *Bombus pullatus* are two closely related Neotropical species with a wide altitudinal and latitudinal distribution that belong to the *Thoracobombus* genus. To the best of our knowledge, there is no genome assembly available for any species of Neotropical *Bombus*. Therefore, the goal of this study is to produce high-quality genomes of *B. pauloensis* and *B. pullatus*. In order to achieve this objective, we obtained long-read sequences using the Oxford Nanopore Technologies platform. We then proceeded to assemble the genomes and annotate these assemblies. As a result, we obtained assemblies of ∼240Mb represented in 72 contigs with an N50 of ∼ 9.08Mb for *Bombus pullatus* and ∼239Mb represented in 66 contigs with an N50 of ∼9Mb for *Bombus pauloensis*. The completeness evaluated by compleasm return a score >99% for both species. It is hoped that these genomes will facilitate a more profound comprehension of the biology of Neotropical bumblebees.

## 1. INTRODUCTION

Bumble bees (*Bombus*) are essential pollinators of a wide range of flowering plants, within both agricultural and natural ecosystems. Despite its importance, is well documented the negative impact of human activities in bumblebees’ richness and abundance in several zones (Goulson et al. 2008; Sánchez-Bayo and Wyckhuys 2019). The subgenus *Thoracobombus* comprises around 50 species from the Old and New World (Williams et al. 2008). All members of this subgenus are characterized by the construction of ground nests, which are typically covered only by herbaceous plant material, such as grass stems (Carder bumble bees) (Williams et al. 2008). In this subgenus, *Bombus pullatus* and *Bombus pauloensis* are two common species of Neotropical ecosystems in Central and South America respectively (Abrahamovich and Díaz 2002). According to Santos Júnior et al. (2022), *B. pauloensis* and *B. pullatus* are closely related species that diverged approximately 5 million years ago following the uplift of the Andes. This geological event likely contributed to their current distinct distribution patterns. Unlike other species, colonies of *B. pauloensis* and *B. pullatus* tend to be larger, often reaching 400 workers, and perennial (Cameron and Williams 2003; Hines et al. 2007; Riaño-Jiménez et al. 2020). Although these species are not considered vulnerable or endangered, is possible that their populations may be declining due the constant human transformation of their habitats.

*Bombus pauloensis* is native to South American ecosystems, with a wide distribution across several countries, including Argentina Bolivia, Brazil, Colombia, Paraguay, Peru, Uruguay, and Venezuela, (Abrahamovich and Díaz 2002; Pinilla-Gallego et al. 2017). While it can be found at elevations ranging from 150 to 3500 meters above sea level (masl), it is most observed between 1800 and 2800 masl (Abrahamovich and Díaz 2002; Pinilla-Gallego et al. 2017). This species exhibits a wide range of color variations, from melanic (completely black) to flavinic (yellow bands on thorax and abdomen) and ferruginous (reddish bands on terga IV-VI) (Ospina T. et al. 1987; Pinilla-Gallego et al. 2017). *Bombus pauloensis* is the most studied South American bumble bee, with research encompassing both basic aspects (biology and ecology) and applications (breeding and crop pollination) (da Silva-Matos and Garofalo 1995; Cameron and Jost 1998; da Silva-Matos and Garófalo 2000; Gonzalez et al. 2004; Gamboa et al. 2015; Riaño-Jiménez et al. 2020; Salvarrey et al. 2020; Cruz et al. 2024). It is a common species that can thrive in a variety of habitats, including highly disturbed environments such as urban gardens, parks, and pastures used for cattle. A remarkable biological characteristic of *B. pauloensis* is its reproductive plasticity. Colonies can exhibit various social structures, including monogyny, polygyny, and competition, which allows for perennial colonies that can last for years, unlike most bumble bee species that have annual colonies (da Silva-Matos and Garofalo 1995; Cameron and Jost 1998; da Silva-Matos and Garófalo 2000). Furthermore, *B. pauloensis* is a highly effective pollinator in high Andean ecosystems and agroecosystems. This has led to the development of industrial captive rearing and commercialization of colonies for crop pollination purposes in several countries, such as Argentina (Almanza et al. 2005; Riaño J. et al. 2015; Salvarrey et al. 2020; Nery et al. 2024). However, little is known about the genome structure and the genetic basis of their unique characteristics.

*Bombus pullatus* is a little-known species recorded from lowlands of Central and Northern South America (Colombia, Costa Rica, Guatemala, Honduras, Nicaragua, Panama, and Venezuela) (Abrahamovich and Díaz 2002). Although it exhibits a wide altitudinal distribution (0 to 3900 masl), *B. pullatus* is most found between 0 and 800 masl (Abrahamovich and Díaz 2002). *Bombus pullatus* is characterized by its melanic coloration, with short, dense black hairs and dark wings (Pinilla-Gallego et al. 2017). Little is known about the biology and ecology of *B. pullatus,* having describing nest architecture and foraging activity in Costa Rica (Janzen 1971; Chavarria 1996; Hines et al. 2007). Nests of *B. pullatus* are like those described in other Thoracobombus such as *B. pauloensis* and *B. transversalis,* constructed over soil, mostly with small cut pieces of dried grass (Hines et al. 2007).

Genetic studies are an essential tool for understanding the vulnerability of bumblebee populations or species to factors linked to their decline, including climatic change, habitat lost, parasites and pathogens, pesticides use and microplastics (Tsvetkov et al. 2021; Boeing et al. 2024; Eldem et al. 2025). Several bumble bee genomes have been sequenced, including those of *B. dahlbomii* (Martínez et al. 2024), *B. huntii* (Koch et al. 2024), *B. impatiens* (Sadd et al. 2015) and *Bombus terrestris* (Sadd et al. 2015). However, the genetic diversity of Neotropical bumble bee species remains largely unknown, as, to the best of our knowledge, no genomes from this region have been sequenced to date.

Therefore, the goal of this study is to generate high-quality genomes of *B. pauloensis* and *B. pullatus*. These genomes represent the first sequenced genomes of Neotropical bumble bees. Given the ecological and biological significance of bumble bees, particularly as key pollinators in Neotropical ecosystems and agroecosystems (Abrahamovich and Díaz 2002), the sequencing of these genomes will enable the identification of gene families involved in detoxification, adaptation, and metabolic processes. This will serve as a valuable tool for understanding the impacts of pesticides, global warming, and diet on bumble bee health. Moreover, this endeavor will contribute to the advancement of knowledge in the fields of evolutionary dynamics and plant-pollinator interactions (Clare et al. 2013), with potential applications in fields such as conservation, evolutionary biology, and agricultural research.

## 2. MATERIALS AND METHODS

### 2.1 Sample Collection and Processing

*Bombus pauloensis* specimens (12 workers) were obtained from three colonies reared in captivity from queens from the municipality of Sopo, Cundinamarca, Colombia (4.908, −73.944). *Bombus pullatus* specimens (12 males) were obtained from a wild colony located in the municipality of Nilo, Cundinamarca, Colombia (4.34920, −74.65430).

The specimens were transported in a living state to the Sequencing Core Facility - GenCore (Universidad de los Andes, Bogotá, Colombia), and subsequently frozen in a refrigerator at −5 °C for 20 minutes. Then, the specimens were dissected in PBS buffer, removing the brain and thoracic muscle tissue (Farris et al. 1999). The tissues were immediately processed for DNA extraction to avoid DNA degradation during storage.

### 2.2 DNA Extraction and Sequencing

Three pools per species were processed: two pools of six brains and one pool of two thoraxes. DNA extraction was performed with a modified protocol of the QIAGEN DNeasy Blood & Tissue Kit. Briefly, each pool was ground using a sterile pestle. Next, each ground tissue was homogenized with 600 µL of Lysis mix (540 µL Buffer ATL and 40 µL Proteinase K). Then, lysis was performed with the following cycles: 5 min disruption at 800 rpm, 25 min incubation at 56°C, 5 min disruption at 800 rpm, 45 min incubation at 56°C with 5 sec vortexing every 10 min, and a final 10 sec final vortexing. The disruption stepes were performed in a BeadBlaster™ 96 Ball Mill Homogenizer. After, 600 µL Buffer AL and 600 µL absolute ethanol were added to each pool. Next, the solution was transferred to a column in three consecutive transfers. The washing steps were performed as specified by the manufacturer. Finally, the elution was performed with 100 µL of DNAse-Free water preheated at 37°C. All columns per each species were mixed during the elution. The concentration of the eluted DNA was verified using Qubit 4.0 with the Qubit dsDNA HS Assay Kit.

Nanopore sequencing libraries were prepared according to the genomic DNA Ligation Sequencing Kit V14 (SQK-LSK114) protocol of Oxford Nanopore Technologies (ONT). Prepared libraries were loaded on PromethION flow cells (R10.4.1) and sequenced with the PromethION 2 (P2) solo device. Finally, basecalling of raw ONT signal data was completed using Dorado v0.7.3 (https://github.com/nanoporetech/dorado) with sup model version 5.0.0.

### 2.3 Assembly

First, the adapters were trimmed using porechop v0.2.4 with the “*-discard_midle”* option (https://github.com/rrwick/Porechop). Reads were filtered into two datasets: Dataset 1) reads of minimum length of 200pb and mean quality of 20, and Dataset 2) reads of minimum length of 10Kb and mean quality of 20 (Table 1). Dataset 1 was used to estimate genome size with k-mer frequency analysis using KAT v2.4.2 (Mapleson et al. 2017) with k-mer lengths of 21, 27 and 31. Dataset 2 was used to assemble the genome with Flye v2.9.4-b1799 (Kolmogorov et al. 2019) with two polishing iterations and without alternative contigs. Then, the draft assembly was polished with medaka v1.12.1 (https://github.com/nanoporetech/medaka) using the reads of the Dataset 1. Contigs with lengths less than 50 Kb were removed because they were likely assembly artifacts. These artifacts often align to longest contigs, exhibit low sequencing depth, and are probably related to issues introduced by the complex pool of DNA used for assembly. (these contigs represent less than 0.005% of the assembly). After, the assembly was checked for contamination using the NCBI Foreign Contamination Screen (FCS, https://github.com/ncbi/fcs) tool suite using the FCS-GX function on the galaxy platform (https://usegalaxy.org/) with the “Animals (Metazoa) - insects” GX-division. A second verification step was performed in Blobtoolkit v4.3.11 (Challis et al. 2020) using coverage and hit data. Briefly, reads from the Dataset 1 were mapped to the assembly using minimap2 v2.28 (Li 2018) and the mapping data were sorted with SAMtools v1.16.1 (Li et al. 2009). Contigs were searches against the core_nt database (accessed 30 August 2024) with blastn v2.16.0 (Camacho et al. 2009) and against the UniProt reference proteome database (accessed 30 August 2024) with diamond v2.1.9 (Buchfink et al. 2014) following the bloobtools2 manual (https://blobtoolkit.genomehubs.org/). For blast and diamond results, the assignments were made at a genus level. Next, duplicate core genes were identified by compleasm v0.2.6 (Huang and Li 2023) using the lineage Hymenoptera orthoDB v10 database. Duplications of these genes between different contigs were manually checked in Gepard v2.1 (Krumsiek et al. 2007) and contigs identified as alternative haplotype of a longer contig or assembling artifacts were removed (in each assembly two contigs were removed). Then, a round of ntLink v1.10.3 (Coombe et al. 2023) was performed with k-mer size of 40 and window of 500, including the *gap_filling* option, to improve the contiguity of the assembly. An additional polishing step was performed after the gap-filling. Briefly, reads of Dataset 1 were mapped to the assembly with minimap2 to run one round of polishing in racon v1.5.0 (Vaser et al. 2017) with default parameters. Finally, a round of medaka with the reads of the Dataset 1 was performed.

**Table 1.**
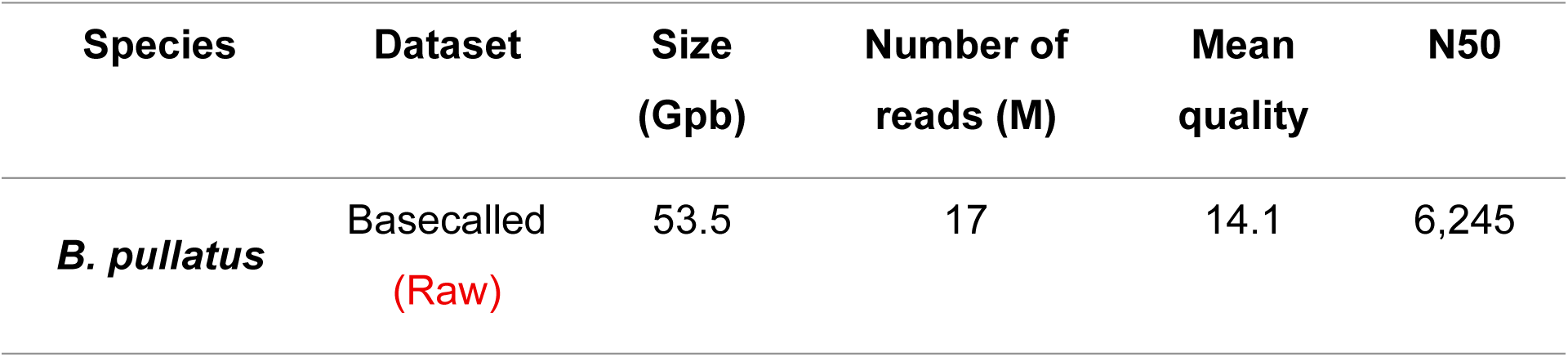
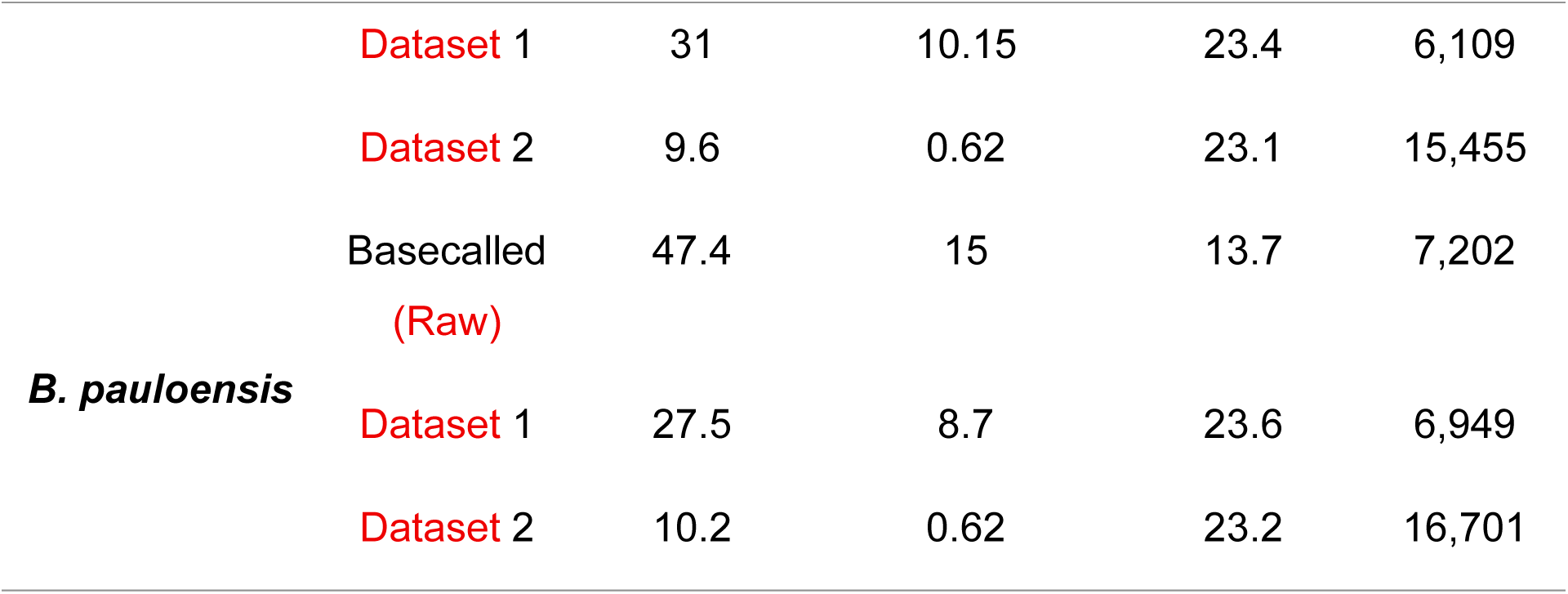
Statistics for Raw and Filtered Reads.

### 2.4 Quality Assement and Species Verification

The quality of the final assembly was checked with Blobtoolkit (Challis et al. 2020) (including coverage data as previously described) that include metrics like number of contigs, N50, depth, GC content and presence/absence of contaminant contigs, and compleasm (Huang and Li 2023) using the Hymenoptera lineage orthoDB v10 database to assess the completeness of the assembly in terms of presence of core single copy ortologues.

In order to verify species identity, three nuclear genes were analyzed: *arginine kinase* (*Argk*), *long-wavelength rhodopsin gene* (*Opsin*), and *phosphoenolpyruvate carboxykinase* (*PEPCK*), these genes had previously been used for phylogenetic analysis (Santos Júnior et al. 2022). The selection of these genes was based on the availability of sequences for both species (*B. pauloensis* and *B. pullatus*) in NCBI. Genomic regions of these genes were identified using BLASTn (Camacho et al. 2009), extracted using Samtools (Li et al. 2009), and, if necessary, the reverse complement was identified using the *revseq* function in Emboss v6.6.0 (Rice et al. 2000). Then, each gene was aligned using MAFFT v7.525 (Katoh and Standley 2013) with the “*--genafpair--maxiterate 1000*” parameters. Finally, phylogenetic reconstruction was performed with *Bombus funerarius* as the outgroup using IQ-TREE2 v2.2.5 with 1,000 UFboostrap replicates (Hoang et al. 2018; Minh et al. 2020). The best-fit evolutionary model of each gene was selected using ModelFinder (Kalyaanamoorthy et al. 2017) under the AICc criteria.

### 2.5 Genome annotation

The identification and masking of transposable elements and repetitive sequences was conducted utilizing the TransposonUltimate pipeline (Riehl et al. 2022). Briefly, transposons were predicted employing the following software: HelitronScanner v1.0 (Xiong et al. 2014), LTRharvest v1.6.2 (Ellinghaus et al. 2008), LTRpred (Drost 2020), MiteFinderII v1.0.006 (Hu et al. 2018), MITE-Tracker v1.0.1 (Crescente et al. 2018), Must v2.4.001 (Ge et al. 2017), NCBI CDD (https://www.ncbi.nlm.nih.gov/Structure/cdd/cdd.shtml), RepeatModeler v2.0.5 (Flynn et al. 2020), RepeatMasker v4.1.6 (https://www.repeatmasker.org/RepeatMasker/), SINE-Finder v1.0.1 (Wenke et al. 2011), Sine-Scan v1.1.2 (Mao and Wang 2017), TIRvish v1.6.2 (Gremme et al. 2013) and TransposonPSI (https://transposonpsi.sourceforge.net/). RepeatMasker was used with the parameters “*-species hymenoptera-e abblast*” against the Dfam database release 3.8 (accessed 17 September 2024). Results of each software were parsed, and duplicates were filtered. Additional transposon copies were identified by searching against software results using CD-HIT v4.8.1 (Fu et al. 2012) and Blastn (Camacho et al. 2009). The final copies were filtered and annotated using RFSB (Riehl et al. 2022). Based on the identified transposons, the genomes were masked using bedtools v2.30.0.

Gene prediction was performed on the masked assembly using BRAKER3 v3.0.6(Gabriel et al. 2024) with the otrhodb v10 database of Hymenoptera as the protein database. Available RNA-seq data from *Bombus* species within the *Thoracobombus* subgenus (*Bombus dahlbomii* [SRR28005379], *Bombus muscorum* [ERR11837462], Bombus pascuorum [SRR6148372], *Bombus pascuorum* [SRR6148369, SRR6148376, SRR6148366], and *Bombus opulentus* [SRR12527964]) were incorporated into the gene prediction process. Then, genes with incomplete models or ORFs shorter than 100 amino acids were filtered out prior to downstream analysis with AGAT v1.4.1 (Dainat et al. 2020).

After, genes were annotated using the Trinotate pipeline (https://github.com/Trinotate/Trinotate/wiki). Briefly, cDNA and protein sequences of each gene model were searched against the swissprot reference databases with BLAST (Camacho et al. 2009). Protein sequences were then analyzed using HMMER v3.4 (hmmer.org) against the Pfam-A database to identify protein domains. Signal peptides were predicted with signalp v6 (Teufel et al. 2022). TMHMM v2.0c (Krogh et al. 2001) was used to identify putative transmembrane regions, and eggnog-mapper v2.1.8 (Cantalapiedra et al. 2021) with the eggNOG database v5.0.2 (Huerta-Cepas et al. 2019) was used for orthology prediction.

Finally, we identified tRNA using tRNAscan-SE v 2.0.12 (Chan et al. 2021) with the default parameters. For others non-coding RNA, cmscan function of infernal v1.1.5 (Nawrocki and Eddy 2013) was used against the Rfam database (accessed 27 September 2024) using the “--rfam--cut_ga--nohmmonly” parameters. Then, results were parsed and summarized.

### 2.7 Whole genome comparison

The available *Bombus pascuorum* genome assembly (GCF_905332965.1) was utilized to identify contigs associated with known chromosomes. Synteny analysis was performed using NGSEP v5.0.0 (Tello et al. 2023) with the *B. pascuorum* genome and annotation as reference. A synteny graph was generated using Rideogram (Hao et al. 2020) to visualize synteny between *B. pascuorum* chromosomes and corresponding contigs in our assemblies. Additionally, paired comparisons were performed between our assemblies and assemblies of other *Thoracobombus* species, focusing on contigs/scaffolds associated with known chromosomes (*B. pascuorum* [GCF_905332965.1], *B. opulentus* [GCA_034509555.1], *B. muscorum* [GCA_963971185.1], and B. *dalhbomii* [GCA_037178635.1]). Dot plots were generated using D-GENIES v1.5.0 (Cabanettes and Klopp 2018) with minimap2 as mapping tools and the “Many repeats” option. For dotplot’s visualization matches were sorted, the short matches were filtered, and the “strong precision” option was enabled.

## 3. RESULTS

High-quality ONT sequencing data provided sufficient coverage to facilitate the successful de novo assembly of the genomes of two *Bombus* species (Table 1). K-mer analysis estimated genome sizes between 239 to 246 Mb and 234 to 241 Mb for *B. pullatus* and *B. pauloensis*, respectively. The final assembly for *B. pullatus* comprised 72 contigs with an estimated genome size of approximately 240 Mb, while the *B. pauloensis* assembly consisted of 66 contigs and an estimated genome size of 239 Mb (Fig. 1A-B). The mean depth of each contig for the second dataset (used for polishing) was approximately 88X for *B. pullatus* and 91X *B. pauloensis* (Fig 1C-D). The N50 for each assembly was approximately 9Mb for both species and the completeness evaluated by compleasm was higher than 99%. Additionally, no contamination was detected in either genome assembly using the FCS and BlobToolKit pipelines. The genomes were verified using three nuclear genes and phylogenetically are grouped with previously known sequences of *B. pauloensis* and *B. pullatus* (Fig. 2).

**Figure 1.**
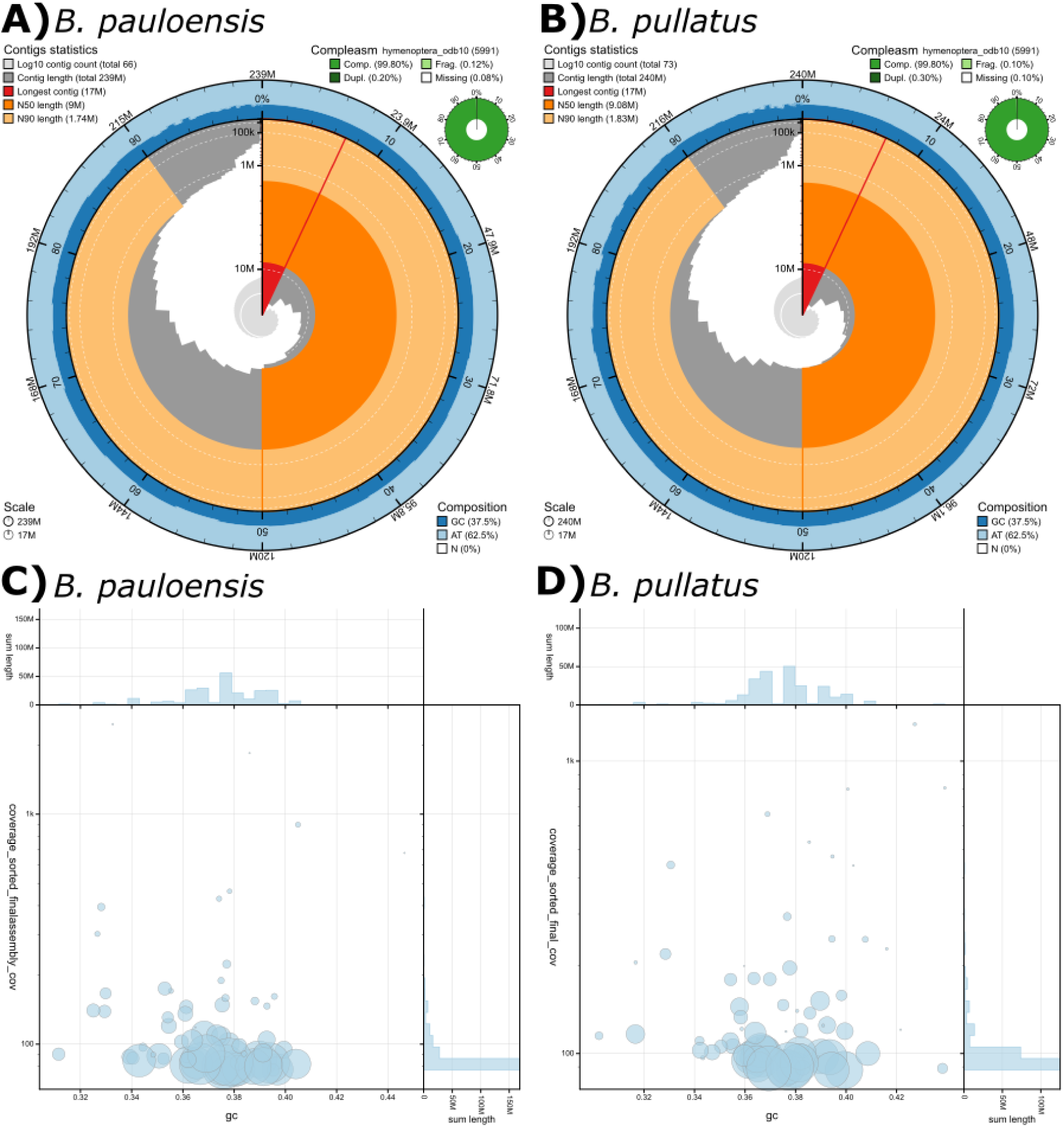
Snail plot visualization of genome assembly statistics for **A)** *B. pauloensis* and **B)** *B. pullatus*. Blob plots showing read depth and gc content of each contig fo **C)** *B. pauloensis* and **D)** *B. pullatus*.

**Figure 2.**
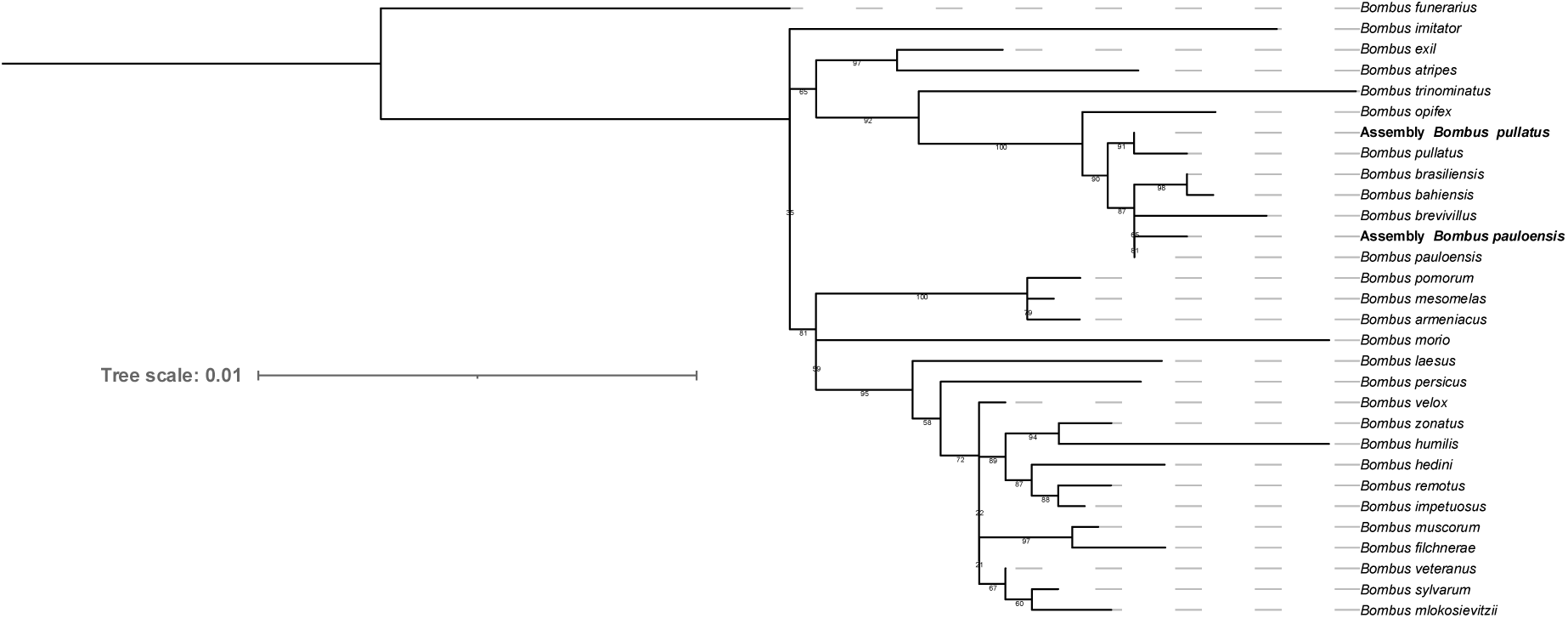
Phylogenetic tree inferred from a concatenated set of genes including *Argk* (521 bp), *Opsin* (648 pb), and *PEPCK* (490 pb) of 27 *Thoracobombus* species. Maximum-likelihood analysis with 1000 ultrafast bootstrap performed in IQ-Tree. *B. funerarius* was used as the outgroup. Support values are shown below branches and represent bootstrap values.

Transposable elements (TEs) analysis shows that approximately 15% of each genome is composed of TEs. The Zator superfamily of DNA transposons exhibited the highest copy number in both assemblies (Table 2). However, the hAT superfamily of DNA transposons has the highest representation in both genomes in terms of base pairs represented by these TEs (Table 2), accounting for approximately 4.2-4.5% of the genome in both species.

**Table 2.** Distribution of TE superfamilies.

| Species | Class | Superfamily | copy number | base pair representation (pb) |
| --- | --- | --- | --- | --- |
| ***Bombus pauloensis*** | **Retrotransposon** | Copia | 187 | 849643 |
|  |  | Gypsy | 1190 | 3103272 |
|  |  | ERV | 2 | 2870 |
|  |  | LINE | 103 | 914918 |
|  |  | SINE | 0 | 0 |
|  | **DNA Transposon** | Tc1-Mariner | 2508 | 8821212 |
|  |  | hAT | 3518 | 9988851 |
|  |  | CMC | 1290 | 9581690 |
|  |  | Sola | 860 | 333438 |
|  |  | Zator | 4603 | 1406201 |
|  |  | Novosib | 43 | 275165 |
|  |  | Helitron | 25 | 162339 |
|  |  | MITE | 1466 | 898266 |
| ***Bombus pullatus*** | **Retrotransposon** | Copia | 221 | 443799 |
|  |  | Gypsy | 1264 | 4253208 |
|  |  | ERV | 3 | 1053 |
|  |  | LINE | 168 | 1188672 |
|  |  | SINE | 0 | 0 |
|  | **DNA Transposon** | Tc1-Mariner | 2383 | 8835882 |
|  |  | hAT | 3687 | 10915817 |
|  |  | CMC | 1393 | 8400228 |
|  |  | Sola | 827 | 272705 |
|  |  | Zator | 4424 | 1260866 |
|  |  | Novosib | 57 | 252602 |
|  |  | Helitron | 47 | 205034 |
|  |  | MITE | 814 | 479891 |

Protein-coding gene annotation in *B. pauloensis* resulted in 10,662 genes with 14,941 transcripts, while *B. pullatus* had 10,681 genes with 14,890 transcripts (Table 3). The average gene length was 8,798 bp in *B. pauloensis* (range: 306-753,695 bp) and 8,704 bp in *B. pullatus* (range: 306-759,035 bp). Both species exhibited an average of 1.4 transcripts per gene. During the annotation, 1987 and 1985 genes in *B. pullatus* and *B. pauloensis,* respectively, could not be annotated, lacking associations with known functions, domains, biological processes, or cellular localizations.

**Table 3.**
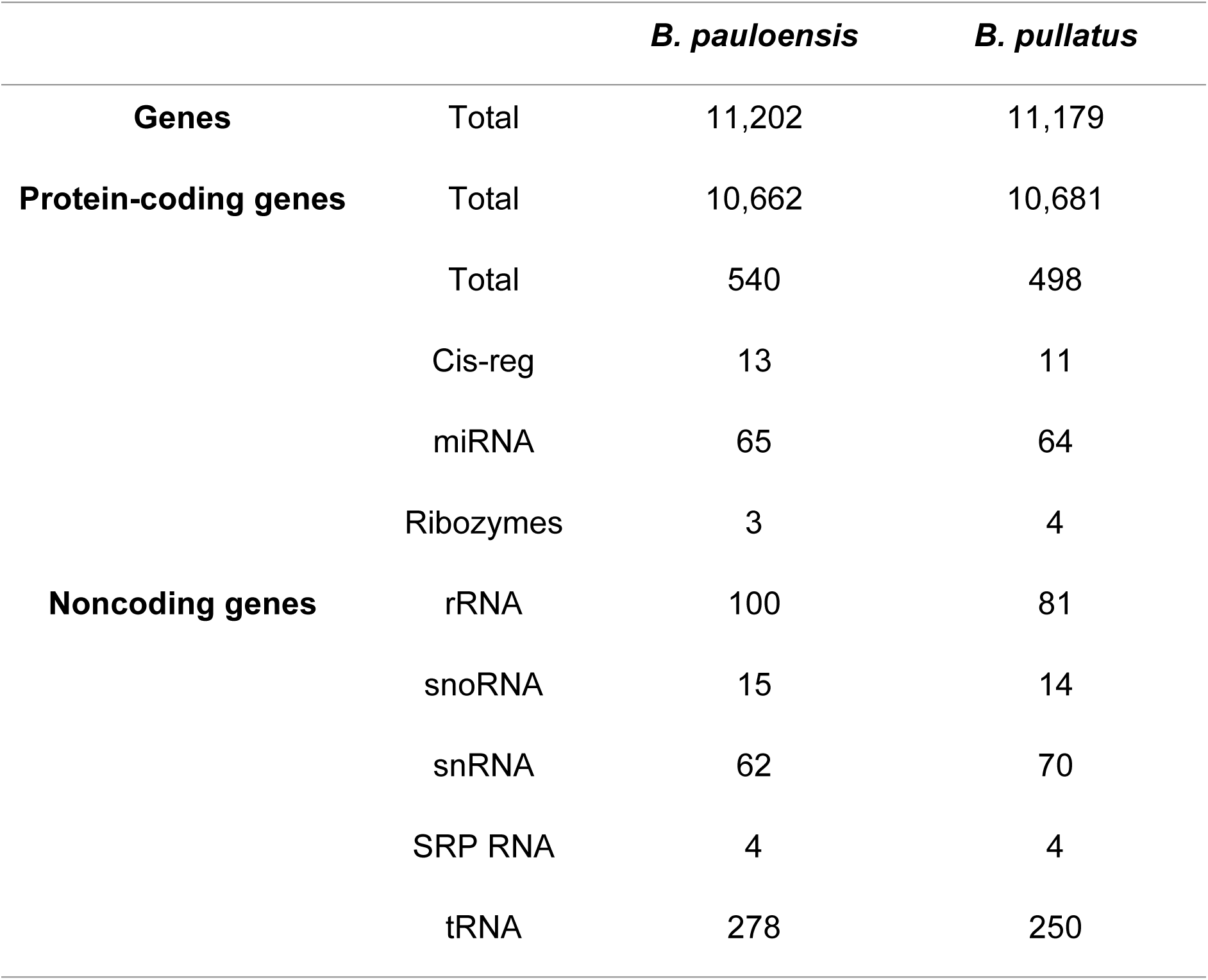
Annotation statistics of assemblies.

Synteny analysis revealed that 45 and 47 contigs in *B. pauloensis* and *B. pullatus*, respectively, could be associated with known chromosomes of *B. pascuorum* (Fig. 3), representing approximately 96% of each genome. This analysis indicated high synteny between our assemblies and the *B. pascuorum* assembly, with evidence of few translocation and inversion events (Fig. 3).

**Figure 3.**
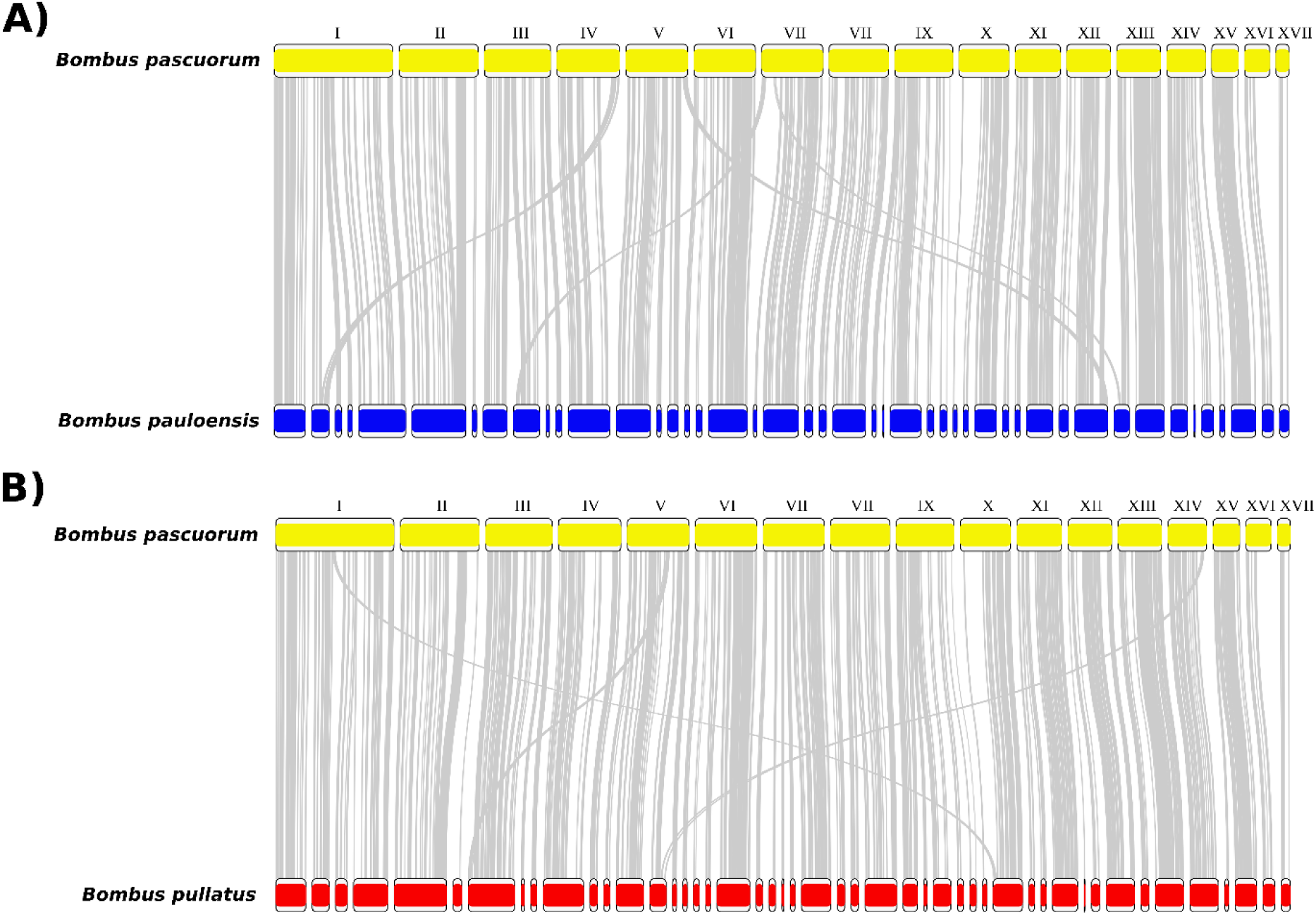
Synteny plot of *B. pascuorum* with **A)** *B. pauloensis* and **B)** *B. pullatus*. Roman numerals represent the chromosomes number for *B. pascuorum*.

Comparison of chromosome-associated contigs between *B. pauloensis* and *B. pullatus* revealed high synteny with limited inversions and translocations (Fig. 4A). Additionally, comparisons with previously reported assemblies of other *Thoracobombus* species revealed some deletions in our assemblies, likely due to the smaller genome sizes of these species (Fig. 4B-I). Furthermore, our assemblies exhibited fewer inversions compared to the *B. opulentus* assembly (Fig. 4D, G).

**Figure 4.**
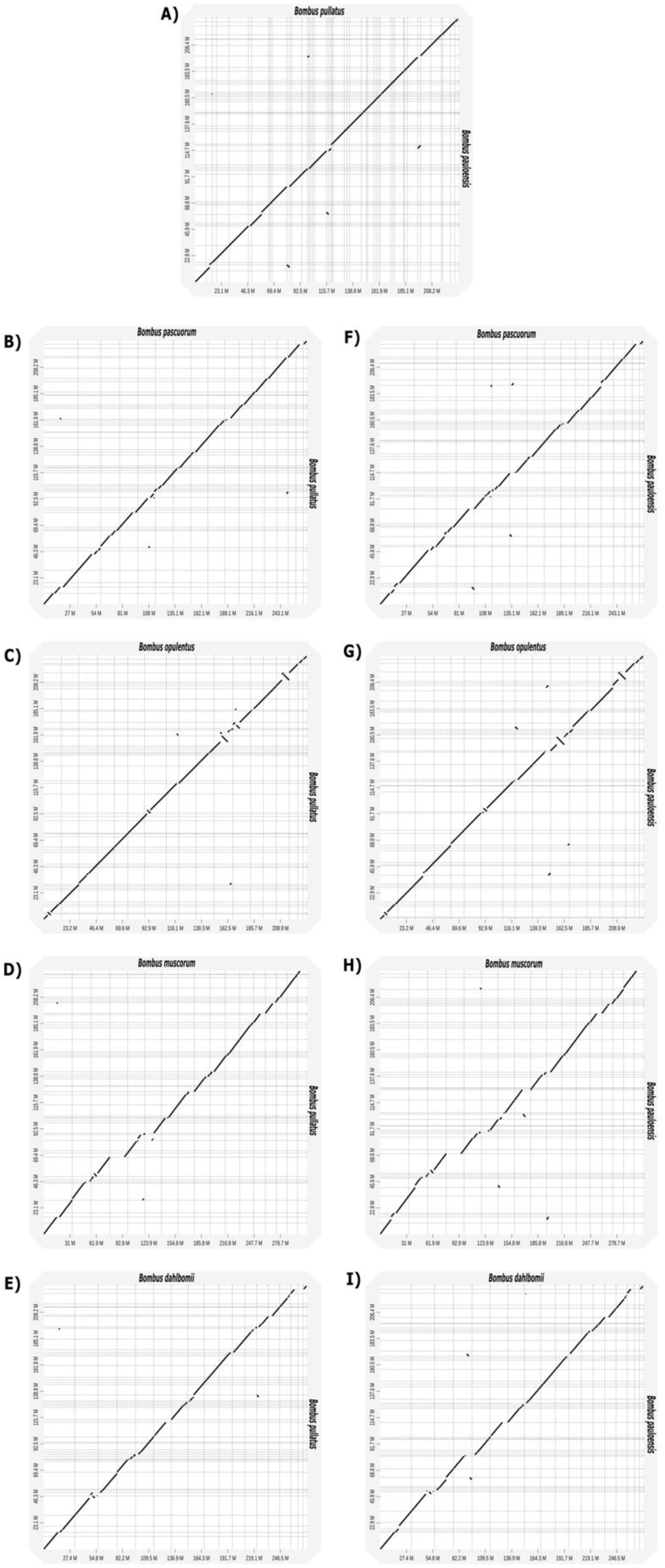
Dot plot of *B. pullatus* against **A)** B. pauloensis, **B)** B. pascuorum, **C)** B. opulentus, **D)** B. muscorum, and **E)** B. dalhbomii. Dotplos of *B. pauloensis* against **F)** B. pascuorum, **G)** B. opulentus, **H)** B. muscorum, and **I)** B. dalhbomii.

## 4. DISCUSSION

In this study, we present the first genome assembly of *B. pauloensis* and *B. pullatus,* two Neotropical species present in Colombia. The genome sizes obtained for *B. pullatus* (240 Mb) and for *B. pauloensis* (239 Mb) are consistent with previously estimated genome sizes for *Bombus* species, which range from 230 Mb to 393 Mb (Sun et al. 2021). These estimates also fall within the range reported for other species within the *Thoracobombus* subgenus (234 Mb to 317 Mb). As expected, the genome sizes of *B. pauloensis* and *B. pullatus* are similar, given their close phylogenetic relationship (Santos Júnior et al. 2022). The results of the k-mer analysis closely predicted the expected genome size for both species, as previously reported in other studies (Koch et al. 2023). With regard to TEs, the proportion of the genome occupied by TEs falls within the expected range for the genus *Bombus* (9-17%) as reported by Sun et al. (2021).

During our analysis, we observed that chromosome naming in previously published genome assemblies of species within the *Thoracobombus* subgenus was often based on scaffold length (Crowley et al. 2023; Broad and Barnes 2024; Martínez et al. 2024), leading to inconsistencies and hindering comparative analyses. For example, it could become challenging to identify homologs of important genes in specific chromosomes across different assemblies due to the varying chromosome order and nomenclature. To address this, we decided to organize and associate our contigs with known chromosomes in *B. pascuorum*, as it is the only species in this subgenus with a reference genome in the RefSeq database (GCF_905332965.1) (Crowley et al. 2023). During these structural comparisons, we identified a few chromosomal rearrangements within the *Thoracobombus* subgenus (Fig. 4). These results suggest a high degree of chromosomal conservation within the *Thoracobombus* subgenus.

The number of protein-coding genes identified in our study falls within the previously reported range for *Bombus* species, which is between 10,000 and 17,000 genes (Heraghty et al. 2020; Sun et al. 2021; Koch et al. 2023; Koch et al. 2024; Martínez et al. 2024). However, the relatively lower number of genes identified in the present study compared to previous ones (Heraghty et al. 2020; Koch et al. 2023; Koch et al. 2024; Martínez et al. 2024) may be attributed to the lack of identification of long non-coding RNAs. Long non-coding RNAs represent the larger group of non-coding genes in *Bombus (Sun et al. 2021)*. It is therefore recommended that future studies concentrate on the identification of long non-coding RNAs in these species of *Bombus*, with a view to achieving a more comprehensive understanding of their gene repertoire and their potential roles the biology of bumble bees.

## 5. CONCLUSION

In conclusion, the present study presents the first high-quality genome assemblies of two closely related neotropical *Bombus* species in Colombia. These genomes exhibit a high degree of contiguity; however, they have not yet been delineated at the chromosome level, a characteristic that distinguishes them from some previously described genomes within this genus. It is evident that further efforts are required in order to complete these assemblies and to improve genome annotation. This will facilitate a more profound comprehension of insecticide resistance, evolutionary dynamics, and adaptations in these and other Neotropical species. These assemblies will be important for identifying genes associated with various traits of interest in these species and ultimately contribute to their conservation. Finally, we hope that this work represents only the first step towards the genetic conservation of Neotropical bumble bees.

## 6. DATA AVAILABILITY

The sequencing data was deposited under the BioProject PRJNA1242843. The reads were deposited in SRA under accession number SRR32927162 and SRR32927163. The genome assemblies were deposited in GenBank under the accession numbers JBNQWY000000000 and JBNQWZ000000000. The scripts used in this study, along with select results pertaining to transposon identification, gene prediction and annotation are deposited in a GitHub repository (https://github.com/andres2901/NeotropicalBumbleBeesAssemblies).

## ACKNOWLEDGMENTS

To Constanza Mendoza, manager of the Mana Dulce Nature Reserve, Nilo Cundinamarca, for allowing the collection of *B. pullatus* samples.

The present study would not have been possible without the sequencing and bioinformatics analysis support provided by the Sequencing Core Facility - GenCore at the Universidad de Los Andes. The High-Performance Computing Service at Universidad de Los Andes is also acknowledged for providing HPC resources necessary to obtain these results in this study. Furthermore, gratitude is extended to the National Environmental Licensing Authority (ANLA) for the approval of the Marco permit for sampling (contract 02492).

## 8. AUTHOR CONTRIBUTIONS

Andres Felipe Lizcano-Salas: Design, data collection, DNA extraction, sequencing, bioinformatic analysis, and wrote the manuscript.

Jesus Jacome-Garcia: Design, *Bombus pauloensis* rearing, data collection, and wrote the manuscript.

Diego Riaño-Jiménez: Design, *Bombus pauloensis* rearing, data collection, wrote the manuscript, and project administration.

Marcela Guevara-Suarez: Design and wrote the manuscript.

